# Different Whole-Brain Functional Connectivity Correlates of Reactive-Proactive Aggression and Callous-Unemotional Traits in Disruptive Children and Adolescents

**DOI:** 10.1101/599548

**Authors:** Julia E Werhahn, Susanna Mohl, David Willinger, Lukasz Smigielski, Alexander Roth, Jilly Naaijen, Leandra M Mulder, Jeffrey C Glennon, Pieter J Hoekstra, Andrea Dietrich, Renee Kleine Deters, Pascal M Aggensteiner, Nathalie E Holz, Sarah Baumeister, Tobias Banaschewski, Melanie C Saam, Ulrike M E Schulze, David J Lythgoe, Arjun Sethi, Michael Craig, Mathilde Mastroianni, Ilyas Sagar-Ouriaghli, Paramala J Santosh, Mireia Rosa, Nuria Bargallo, Josefina Castro-Fornieles, Celso Arango, Maria J Penzol, Marcel P Zwiers, Barbara Franke, Jan K Buitelaar, Susanne Walitza, Daniel Brandeis

## Abstract

**Background:** Disruptive behavior in children and adolescents can manifest itself in reactive (RA) and proactive (PA) aggression and is modulated by callous-unemotional (CU) traits and comorbidity. Research on aggression subtype-specific neural correlates is limited and the role of comorbid symptoms largely neglected.

**Methods:** The current multi-center study extended previous efforts by investigating unrestricted resting state functional connectivity (rsFC) alterations. The large sample (n = 207) of children and adolescents aged 8 – 18 years (mean age = 13.30 ± 2.60 years) included 118 cases with disruptive behavior (80 diagnosed with Oppositional Defiant Disorder and/or Conduct Disorder) and 89 controls. Attention-deficit/hyperactivity disorder (ADHD) and anxiety symptoms were added as covariates. We measured changes in global and local voxel-to-voxel rsFC using functional magnetic resonance imaging at 3T (mean acquisition time = 8 min 25 sec).

**Results:** Compared to controls, cases demonstrated altered rsFC including frontal areas when anxiety but not ADHD symptoms were considered. Within cases, RA and PA scores related to changes in global and local rsFC in central gyrus and precuneus previously linked to aggression-related impairments. CU trait severity correlated with global rsFC alterations including inferior and middle temporal gyrus implicated in empathy, emotion, and reward-related activity. Importantly, most observed aggression subtype-specific patterns could only be identified when ADHD and anxiety problems were also accounted for.

**Conclusions:** The current study clarifies that distinct though overlapping brain connectivity measures can disentangle differing manifestations of aggressive behavior. Moreover, our results highlight the importance of considering comorbid symptoms for detecting aggression-related rsFC alterations.

## INTRODUCTION

Severe aggressive behavior in children and adolescents is a main characteristic of DSM-5 aggression-related disorders (American Psychiatric Association, 2013). Conduct disorder (CD) is described as persisting violation of rules, norms, and rights including physical and psychological aggressive behaviors. Oppositional defiant disorder (ODD) identifies a pattern of angry, argumentative, and vindictive behavior. The combination of childhood CD and comorbid attention-deficit/hyperactivity disorder (ADHD) is thought to be associated with persistent antisocial behavior (Moffitt, 1993; Moffitt *et al.*, 2002; Frick, 2016). Besides ADHD (Loeber *et al.*, 2000), frequently exhibited comorbid anxiety levels (Frick, 2012; Frick *et al.*, 2013) seem crucial for subtyping aggression on a psychophysiological level (Fanti, 2016).

On a neural level, frontal, striatal, and limbic functional alterations have been identified in CD and ODD (Blair, Veroude and Buitelaar, 2016). Resting state functional magnetic resonance imaging (rs-fMRI) allows to identify distinct highly functionally connected brain regions active during the absence of goal-directed behaviors (Raichle *et al.*, 2001; Greicius *et al.*, 2003; Fox *et al.*, 2015). Male adolescents with CD compared to controls were shown to exhibit reduced functional activity in limbic areas (Zhou *et al.*, 2015) along with parietal, cerebellar, and temporal areas including the middle temporal gyrus (MTG) (Wu *et al.*, 2017). Further rs-fMRI studies reported increased (Lu, Zhou, Wang, *et al.*, 2017) and reduced resting state functional connectivity (rsFC) (Lu *et al.*, 2015; Broulidakis *et al.*, 2016; Zhou *et al.*, 2016; Lu, Zhou, Wang, *et al.*, 2017) paralleling reduced anisotropy at the structural level in male and female adolescents with CD (González-Madruga *et al.*, 2019). For instance, male adolescents with CD exhibited reduced rsFC in the central gyrus (Lu *et al.*, 2015). The precuneus as part of the default mode network (DMN) has been implicated in increased (Lu, Zhou, Wang, *et al.*, 2017) and decreased rsFC (Zhou *et al.*, 2016; Lu, Zhou, Zhang, *et al.*, 2017). Males compared to females with CD were shown to exhibit decreased spontaneous activity during rest in brain areas including left inferior temporal gyrus (ITG)/MTG and right postcentral gyrus (Cao *et al.*, 2018). Compared to controls, boys with CD and comorbid ADHD demonstrated increased rsFC of DMN seeds with voxel clusters including ITG and MTG (Uytun *et al.*, 2016). These previous rsFC studies, however, are mostly restricted to case-control comparisons in male adolescents with CD and further limited by small sample sizes.

Aggression is a heterogeneous phenotype and has frequently been subdivided into reactive (RA) and proactive (PA) aggressive behavior (Dodge and Coie, 1987; Vitiello and Stoff, 1997). Individuals with RA are prone to perceive behavior of others as provocation or threat and react aggressively (Smeets *et al.*, 2016). In contrast, individuals with PA tend to show planned, instrumental aggressive behaviors. Few MRI studies have addressed the brain correlates of RA and PA in disruptive children and adolescents. There is evidence for RA and PA relating to distinct seed-based rsFC patterns involving precuneal areas (Werhahn *et al.*, 2018).

Callous-unemotional (CU) traits are reported to be an important modifier of aggression. These behaviors characterized by limited prosocial emotions have been added to further specify CD in the DSM-5 (American Psychiatric Association, 2013) and were shown to reduce treatment response and worsen clinical outcome (Wilkinson, Waller and Viding, 2016; Bakker *et al.*, 2017). Moreover, CU traits seem associated with impaired empathy (Blair, Leibenluft and Pine, 2014; Ciucci *et al.*, 2015), neurocognitive dysfunctions in emotion and reward learning processes (Reidy *et al.*, 2017), and adult psychopathy (Frick and White, 2008; Frick *et al.*, 2013). There is evidence for CU traits associated with both RA and PA (Pechorro *et al.*, 2017) or PA only (Urben *et al.*, 2018). Importantly, youths with CU and CU-related traits but different levels of anxiety also differ in behavioral characteristics (Kimonis *et al.*, 2012; Fanti, Demetriou and Kimonis, 2013). On a neural level, rs-fMRI studies on CU traits in children and adolescents are limited. Male adolescents with higher CU and CU-related traits showed altered amygdala subregional rsFC with voxel clusters including parietal, frontal, temporal, and cerebellar areas (Aghajani *et al.*, 2016, 2017). Disruptive children and adolescents showed CU subdimension-specific seed-based rsFC including clusters in central gyrus and precuneus (Werhahn *et al.*, 2018). Other studies reported CU-related rsFC alterations in the DMN (Cohn *et al.*, 2015; Pu *et al.*, 2017; Thijssen and Kiehl, 2017) along with further frontal, temporal, parietal, and cerebellar areas (Espinoza *et al.*, 2018). Only one study in adults addressed anxiety symptoms and reported an interaction effect with psychopathy levels (Motzkin *et al.*, 2011), despite their prominent behavioral and psychophysiological effects (Fanti, Demetriou and Kimonis, 2013; Guelker *et al.*, 2014; Dadds *et al.*, 2017). In sum, knowledge on rsFC correlates of CU traits is limited, and the influence of comorbid anxiety unknown.

The current study was designed to substantially extend prior work by investigating unrestricted graph-theoretical-based whole-brain voxel-to-voxel rsFC in a large sample of male and female children and adolescents with 118 aggressive cases and 89 healthy controls. We examined the rsFC underpinnings of the RA and PA dimensions as well as of CU traits, and took co-occurring ADHD and anxiety symptoms into account. We calculated the Intrinsic Connectivity Contrast (ICC) to measure changes in network centrality (Martuzzi *et al.*, 2011) and the Integrated Local Correlation (ILC) to examine changes in local coherence (Deshpande *et al.*, 2009). We hypothesized to find 1) group differences in rsFC in frontal regions, 2) RA and PA-related rsFC alterations in the precuneus, 3) rsFC alterations for CU traits beyond (para-)limbic regions (Werhahn *et al.*, 2018), and 4) these alterations to be partly modulated by ADHD and anxiety symptoms.

## MATERIALS AND METHODS

### Participants

For the present study, 118 children and adolescents (*n* males = 99) aged 8-18 years (mean age = 13.23, *SD* = 2.68) with a diagnosis of ODD and/or CD and/or clinically relevant aggression scores along with 89 healthy controls were included. Cases were recruited from resident hospitals, ambulatories, and eligible schools at nine sites in Europe in the framework of the joint EU-MATRICS and EU-Aggressotype projects. Clinically relevant aggression was defined as aggression scores in the clinical range (*T* > 70) according to the Child Behavior Checklist (CBCL), Youth Self Report (YSR), or Teacher Report Form (TRF) (Achenbach, 1991). Exclusion criteria for cases were a primary DSM-diagnosis of depression, anxiety, psychosis, or bipolar disorder for cases and for the typically developed comparison group a DSM-diagnosis and a clinically relevant aggression score in the CBCL, TRF, or YSR. Furthermore, participants were excluded in case of contraindications for MRI scanning (i.e., braces, metal implants), insufficient language skills, or an IQ < 80 (Wechsler, 2003). Parents or legal representatives of all participants gave written informed consent. The participating sites obtained ethical approval separately from their local ethics committees. Further information on the study sample is provided in the Supplement.

### Clinical Assessments

To be included, cases had to exhibit a DSM-diagnosis of CD and/or ODD according to the semi-structured interview Kiddie-Schedule for Affective Disorders and Schizophrenia, present and lifetime version (K-SADS-PL) (Kaufman *et al.*, 1997) or a clinically relevant score (*T* > 70) on the aggression or rule-breaking behavior subscale of the CBCL, YSR, or TRF (Achenbach, 1991). The assessment of these clinical questionnaires and the K-SADS-PL in the typically developed comparison group ensured the absence of a DSM-diagnosis or clinically relevant aggression scores. Anxiety symptoms were measured using the YSR, as internalizing problems including anxiety were shown to exhibit higher validity (Leung *et al.*, 2006) and reliability (Gomez, Vance and Gomez, 2014) compared to the CBCL and TRF. The Reactive-Proactive Questionnaire (RPQ) is a self-report on the frequency of RA and PA symptoms (Raine *et al.*, 2006). As previously reported (Cima *et al.*, 2013), RA and PA symptoms within cases were correlated strongly in the current study (*r* = 0.63, *p* < 0.001). CU traits were assessed by the Inventory of Callous-Unemotional traits (ICU) (Essau, Sasagawa and Frick, 2006). Here, we used the parent-reported version of the ICU, as it seems to indicate better CU traits-related variables compared to the self-report (Docherty *et al.*, 2017). To assess comorbid ADHD symptoms, parents answered the SNAP-IV (Swanson, 1992). The inattention and hyperactivity/impulsivity subdomains were used in the present study. Four subscales of the Wechsler Intelligence Scale for Children (Block Design, Similarities, Vocabulary, Picture Completion) (Wechsler, 2003) enabled the estimation of an IQ in order to ensure sufficient intellectual and cognitive functioning for inclusion (IQ ≥ 80). Further tests conducted as part of a larger study protocol are described in the Supporting Information.

### Image Acquisition

MRI scanning took place at nine sites across Europe, with six sites using Siemens 3T scanners, two sites Philips 3T scanners, and one site a GE 3T scanner. Besides a T1-weighted structural scan (see Supplementary Table 1), a T2*-weighted whole-brain echo planar imaging (EPI) resting state sequence (TR 2.45s or less, at least 32 slices) with an average acquisition time of 8 min 25 sec was performed (see Supplementary Table 2). Participants were instructed to lie still, look at a white crosshair presented against a black background, and not to think about anything in particular. More information is provided in the Supporting Information.

### Preprocessing

Functional magnetic resonance imaging (fMRI) data were preprocessed using SPM version 12 (http://www.fil.ion.ucl.ac.uk/spm). After realignment and unwarping (Andersson *et al.*, 2001) followed by slice timing correction, multi-echo images were weighted by their echo time (TE) using Matlab [The MathWorks, MA, USA]. All rs-fMRI data were functionally normalized to the Montreal Neurological Institute template brain shown to reduce variability across subjects (Calhoun *et al.*, 2017) using SPM-based functional connectivity CONN toolbox version 17f (Whitfield-Gabrieli and Nieto-Castanon, 2012) in Matlab. Functional scans were smoothed with a Gaussian 6 mm full width at half-maximum kernel. Functional outliers exceeding > 3 standard deviations from the mean intensity across blood-oxygen-level-dependent (BOLD) time series and > 0.5 mm composite scan-to-scan motion were identified with ART (Artifact Detection Toolbox, www.nitrc.org/projects/artifact_detect) within CONN. We excluded 10 cases from further analysis with functional scans exceeding the threshold of >5% of the highest mean Root Mean Square (RMS) framewise displacement (FD) (cut-off = 0.95 RMS-FD), as applied recently in adolescents with ADHD (von Rhein *et al.*, 2016). Moreover, 23 cases and three controls were not included due to missing (*n* = 2) or insufficient quality of the structural (*n* = 10) or functional (*n* = 14) scans. During denoising, the aCompCor strategy (Behzadi *et al.*, 2007) implemented in CONN enabled physiological and motion-related noise reduction (Whitfield-Gabrieli and Nieto-Castanon, 2012) and improved interpretability of resulting correlation patterns (Chai *et al.*, 2012; Saad *et al.*, 2013). Besides linear detrending, BOLD time series were band-pass filtered between 0.008 Hz and 0.09 Hz (Fox *et al.*, 2005).

### Functional Connectivity Analyses

After case-control group comparisons, we calculated changes in rsFC related to RA and PA symptoms along with CU traits within cases. Intrinsic Connectivity Contrast (ICC) was used to determine the magnitude of degree or network centrality by calculating the global strength of connectivity for each voxel with other voxels in the rest of the brain [Martuzzi *et al.*, 2011]. ICC has been frequently applied in recent rsFC studies (Van Ombergen *et al.*, 2017; Vatansever *et al.*, 2017; Walpola *et al.*, 2017; Browndyke *et al.*, 2018; Cassady *et al.*, 2018). It has the advantage of circumventing the need for a priori definitions of regions of interest (Martuzzi *et al.*, 2011). Integrated Local Correlation (ILC) was used to compute changes in average local connectivity patterns by integrating voxel-wise spatial correlations (Deshpande *et al.*, 2009). Compared to other local coherence approaches, ILC is independent of image resolution and a predefinition of the neighborhood is not necessary. ILC seems to be tissue-specific and widely independent of physiological noise (Deshpande *et al.*, 2009). First-level analysis was used to calculate voxel-to-voxel covariance matrices for each subject, which further served to calculate ICC and ILC. Correlation coefficients from these functional connectivity analyses were z-transformed and therefore normalized. Within second-level analysis, random-effects analysis of covariance was then employed to calculate group comparisons. Linear regressions tested the influence of the distinct aggression subtypes separately. Besides adding site as dummy coded covariate, we adjusted for parent-reported ADHD (Broulidakis *et al.*, 2016) and self-reported (Leung *et al.*, 2006; Gomez, Vance and Gomez, 2014) anxiety levels (Motzkin *et al.*, 2011) given their crucial role shown in behavioral studies on aggression (Fanti, Demetriou and Kimonis, 2013; Guelker *et al.*, 2014; Frick, 2016; Dadds *et al.*, 2017). Additional sensitivity analyses including further covariates such as age and sex are provided in the Supporting Information. For all results, an uncorrected height threshold of *p* < 0.001 with a *p* < 0.05 false-discovery-rate (FDR) cluster threshold was applied.

## RESULTS

### Sample Characteristics

As presented in Table 1, our sample consisted of 118 cases and 89 healthy controls. Forty-eight cases had a diagnosis of ODD, 25 of CD and ODD, and seven of CD. Seventy-seven cases presented with scores in the clinical range (T > 70) on the aggression or the rule-breaking behavior subscale of the CBCL and 41 cases on both subscales. Thirty-eight cases had an aggression score in the clinical range but no DSM-diagnosis. These cases exhibited comparable RA values (*M* = 12.67, *SD* = 5.62) and CU traits (*M* = 33.61, *SD* = 10.01), and lower PA scores (*M* = 3.91, *SD* = 4.00) compared to cases with a diagnosis (RA: *M* = 12.50, *SD* = 4.87; PA: *M* = 5.27, *SD* = 5.37; CU traits: *M* = 33.71, *SD* = 10.25). Most participants presented with comorbid ADHD symptoms (*n* = 103), 66 cases had anxiety problems, and 70 cases received medication. Within cases, only the positive correlation between RA (*r* = 0.32, *p* = 0.004) and anxiety symptoms and between CU traits and ADHD inattention subscale (*r* = 0.32, *p* = 0.001) yielded significance. Further associations are reported in the Supplemental Results.

**Table 1.**
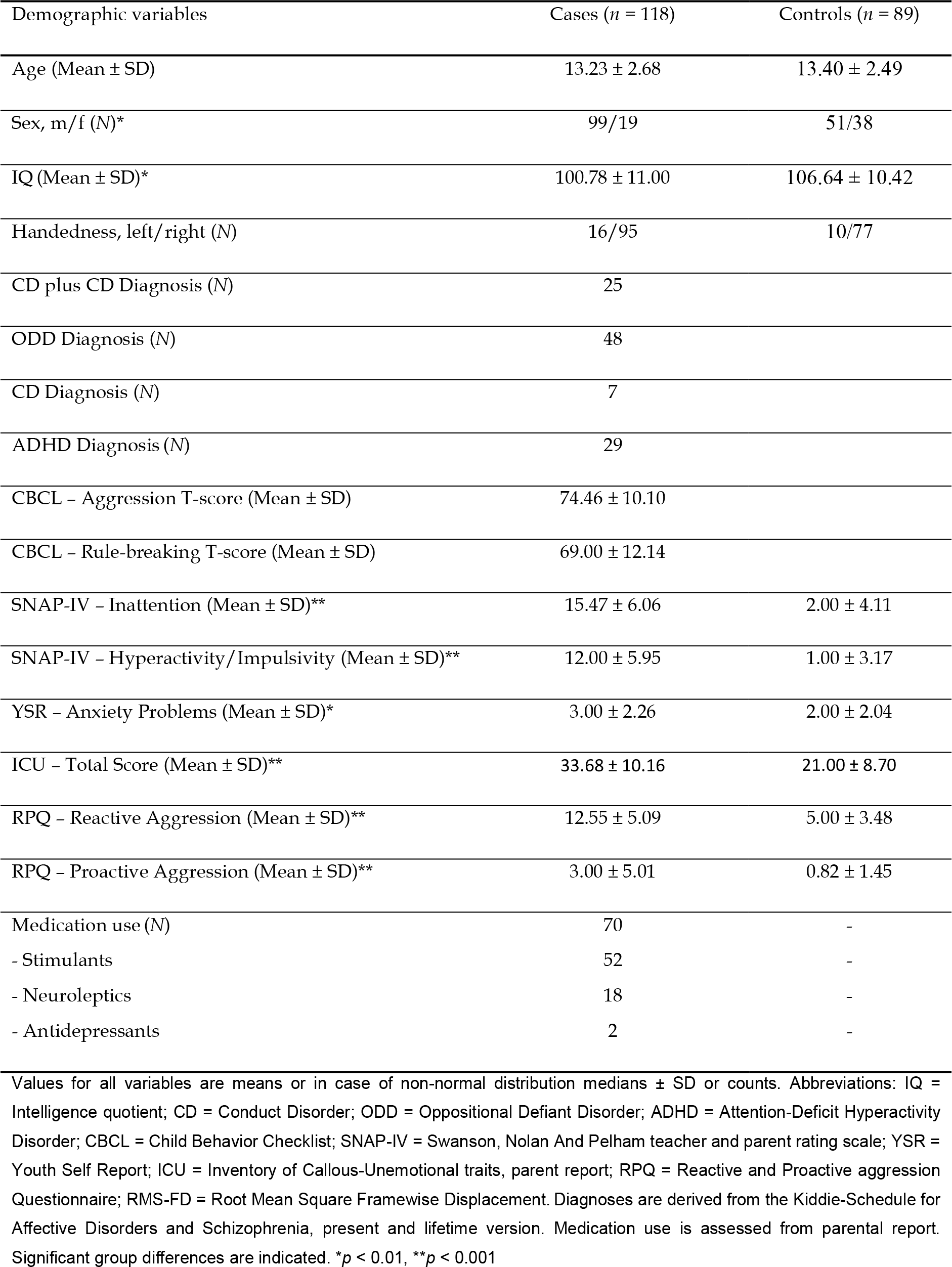
Sample characteristics

### Group Differences in Functional Connectivity

Only after the exclusion of ADHD symptoms as additional covariate, cases showed increased ILC in a cluster including bilateral frontal pole extending to the medial frontal cortex compared to controls (*T*(142) = 6.12, *p*-FDR < 0.05). Moreover, cases demonstrated reduced ICC in a cluster including the right occipital pole (*T*(142) = −5.19, *p*-FDR < 0.05). These group differences survived the control for additional covariates including age and sex.

### Functional Connectivity Correlates of Reactive and Proactive Aggression

Both higher levels of RA and PA related to altered ICC in the left superior parietal lobe and lateral occipital cortex. With higher levels of RA, ICC decreased in these areas in the left hemisphere, while with higher levels of PA, ICC increased in these regions in the right hemisphere. The RA-related patterns extended to the left central gyrus and the PA-related pattern to the precuneus (Fig. 1A, see Supplementary Table 4). Moreover, cases demonstrated decreased ILC with increasing levels of both RA and PA symptoms in comparable left hemispheric voxel clusters including superior parts of the parietal lobe and lateral occipital cortex along with the supramarginal gyrus, extending to angular gyrus. Only the PA symptoms-related patterns extended to left central gyrus (Fig. 1B, see Supplementary Table 5).

**Figure 1.**
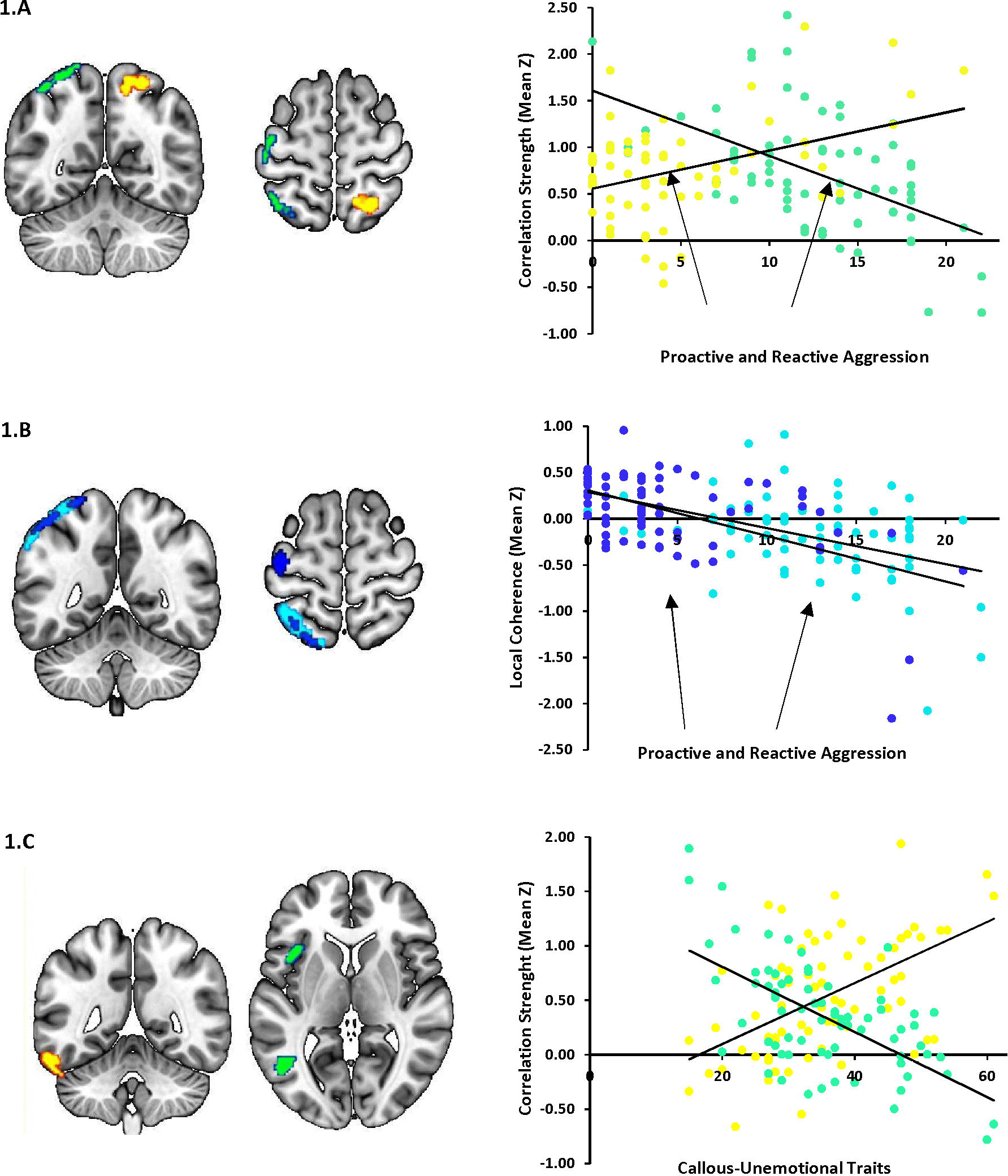
Aggression subtype-specific whole-brain voxel-to-voxel connectivity correlates of reactive and proactive aggression (RA/PA) and callous-unemotional (CU) traits. Higher RA and PA scores related to decreased versus increased intrinsic correlation contrast (ICC) in a cluster including left and right superior parts of parietal lobe and occipital cortex colored in green and yellow, respectively. Moreover, a cluster in left central gyrus exhibited RA-specific decreased ICC and the cluster for PA scores extended to precuneus (I.A). Regarding integrated local correlation (ILC), RA and PA related to decreased ILC colored in light and dark blue, respectively, in left hemispheric areas extending from superior parts of parietal lobe and occipital cortex to angular gyrus. Further, a cluster in left central gyrus exhibited a PA dependent decrease in ILC (I.B). Higher CU traits correlated with increased ICC in left inferior temporal gyrus and decreased ICC in left hemispheric clusters including medial and inferior temporal gyrus and frontal and central opercular regions (I.C). For all aggression subtype-specific associations, scatterplots depict the direction (blue and green colors indicate decreased, yellow color indicates increased), with functional connectivity strength (ICC) or local coherence (ILC) values (y-axis) plotted against scores of RA and PA and CU traits (x-axis). Z-values are Fisher transformed correlation coefficients and averaged across all observed voxels in case of more than one cluster. Statistical threshold of the connectivity results visualized in representative coronal and axial views is *p* < 0.001, with false discovery rate (FDR) cluster-level corrected (*p* < 0.05) for multiple comparisons.

### Functional Connectivity Correlates of Callous-Unemotional Traits

With higher levels of CU traits, ICC increased in a left hemispheric cluster including the ITG and the cerebellum. In contrast, ICC decreased in left hemispheric clusters that extended from the ITG and MTG and inferior lateral occipital cortex to frontal and central opercular areas including the insula lobe (Fig. 1C, see Supplementary Table 4).

## DISCUSSION

The present study advanced previous work by conducting unrestricted whole-brain rsFC analyses in a large sample of children and adolescents. Comorbid anxiety and ADHD symptoms were taken into account. First, only when we did not control for ADHD symptoms, cases showed altered voxel-wise rsFC compared to healthy controls including frontal clusters implicated in cognitive control. Second, both RA and PA correlated with voxel-wise alterations in rsFC including superior parts of the parietal lobe linked to activities like attentional control. Third, our analyses yielded distinct voxel-wise rsFC patterns for RA and PA in left central gyrus and/or precuneus implicated in aggression and cognitive control-related processes. Fourth, distinct CU traits-specific altered ICC included temporal and cerebellar regions suggested to play an important role during fear, moral, and reward.

Importantly, most aggression subtype-specific rsFC findings depended on the additional control of both ADHD and anxiety symptoms, suggesting that these symptoms have suppressor effects, which can obscure aggression subtype-specific patterns. This underlines the importance of considering both ADHD (Broulidakis *et al.*, 2016; Uytun *et al.*, 2016) and anxiety (Motzkin *et al.*, 2011). Within cases, we found mostly small and insignificant behavioral associations of RA and PA symptoms and CU traits with ADHD and anxiety levels, which might further point to heterogeneous underlying profiles within these manifestations of aggression. The crucial role of ADHD and anxiety regarding subtyping aggression is also corroborated at the psychophysiological level (Fanti, 2016). Behavioral studies suggest a correlation between anxiety symptoms and RA and PA (Fite *et al.*, 2014; Fung *et al.*, 2015) or RA symptoms particularly (Vitaro, Brendgen and Tremblay, 2002; Marsee, Weems and Taylor, 2008) as in our sample. RA has been related to anxiety and ADHD problems (Smeets *et al.*, 2016) and internalizing symptoms (Fite *et al.*, 2014), and therapeutic approaches address anxiety to treat RA symptoms (Blair, Leibenluft and Pine, 2014). CU traits seem typically associated with reduced anxiety (Frick *et al.*, 2013; Eisenbarth *et al.*, 2016). Cases with CU traits and co-occurring high levels of anxiety might represent a distinct phenotype with different clinical outcomes (Fanti, Demetriou and Kimonis, 2013) and neural correlates (Sethi *et al.*, 2018). To conclude, our results emphasize to differentiate between aggression subtypes and consider comorbid anxiety and ADHD symptoms.

In line with previous efforts, cases exhibited rsFC alterations in the frontal pole compared to controls. Importantly, we only found this cluster that extended to the medial frontal cortex implicated in cognitive control (Ridderinkhof *et al.*, 2004) when we considered anxiety but not ADHD symptoms. The importance of taking anxiety levels into account is in line with task-based fMRI reports in antisocial youths (Byrd, Loeber and Pardini, 2014), while the role of ADHD could not be supported (Broulidakis *et al.*, 2016; Werhahn *et al.*, 2018).

Our RA and PA-related analyses yielded decreased ICC in clusters including superior parts of the parietal lobe and lateral occipital cortex, however, in different hemispheres. Decreased ILC included the angular gyrus implicated in altered rsFC of individuals with antisocial personality disorder (Tang *et al.*, 2016) and CU-related traits (Espinoza *et al.*, 2018). Activity of the angular gyrus as part of the DMN (Andrews-Hanna, Smallwood and Spreng, 2014) has been associated with CU-related traits during emotional processing (Anderson *et al.*, 2017) and with moral behaviors (Raine and Yang, 2006; Fumagalli and Priori, 2012; Pujol *et al.*, 2012; Boccia *et al.*, 2017). The lateral occipital cortex has been implicated in reduced functional activity during rest in adolescents with CD (Haney-Caron, Caprihan and Stevens, 2014). The superior parietal lobe has been linked to attentional control (Pessoa, Kastner and Ungerleider, 2002), learning from reward and punishment (Finger *et al.*, 2011), and fearful expressions (Marsh *et al.*, 2008; Peraza *et al.*, 2015). In sum, our results suggest overlapping rsFC alterations related to both RA and PA symptomatology in brain regions involved in attention, decision-making, emotions, and CU-related traits.

As expected and in line with behavioral studies (Marsee and Frick, 2007; Fite *et al.*, 2008, 2010; Marsee *et al.*, 2011) and previous work, we found distinct rsFC patterns associated with RA and PA symptoms. The precuneus and central gyrus have been associated with different forms of aggressive behavior and cognitive control-related processes known to be impaired in disruptive behavior (Blair, Veroude and Buitelaar, 2016). Interestingly, SN seed-based rsFC with clusters including these regions related to CU subdimensions (Werhahn *et al.*, 2018). Firstly, and as expected, altered voxel-wise rsFC in the precuneus related to PA symptoms. Aberrant rsFC of the precuneus has been linked to impulsivity (Lu, Zhou, Wang, *et al.*, 2017), CU-related traits (Aghajani *et al.*, 2016), ADHD and severe temper outbursts (Roy *et al.*, 2018). During rest, the precuneus showed altered functional activity in male adolescents with CD (Zhou *et al.*, 2016) and rsFC in male juvenile offenders (Chen *et al.*, 2015) and antisocial (Tang, Jiang, *et al.*, 2013; Tang *et al.*, 2016) and psychopathic adults (Motzkin *et al.*, 2011; Philippi *et al.*, 2015). The precuneus as part of the DMN (Greicius *et al.*, 2003) has been linked to self-reflection (Cavanna, 2007) and moral reasoning (Boccia *et al.*, 2017). It seems to be involved in cognitive control and aggressive interactions (Fanning *et al.*, 2017). Secondly, the left central gyrus showed decreased ICC with higher PA and decreased ILC with higher RA scores. Altered precentral gyrus rsFC has been reported in psychopathic adults (Tang *et al.*, 2016; Espinoza *et al.*, 2018) and decreased postcentral gyrus rsFC in adolescents with CD (Lu *et al.*, 2015). Despite these neural findings and the behavioral correlation between CU traits and both RA and PA symptoms (Kimonis *et al.*, 2008; Feilhauer, Cima and Arntz, 2012; Pechorro *et al.*, 2017), RA and PA-related rsFC did not overlap with CU traits-specific patterns.

CU traits related to voxel-wise rsFC alterations in left hemispheric ITG, MTG, and opercular regions along with the cerebellum linked to emotion, reward, and moral-related processes. Previous rs-fMRI studies in male adolescents reported CU-related rsFC alterations in MTG and ITG (Aghajani *et al.*, 2016; Thijssen and Kiehl, 2017), and CD-related diminished right MTG activity (Wu *et al.*, 2017). MTG activity has been reduced also in CD (Cao *et al.*, 2018) and during a risk-taking task (Dalwani *et al.*, 2014). The MTG exhibited the highest discriminate power within the DMN to distinguish individuals with antisocial personality disorder from controls and the cerebellum enabled the best differentiation overall (Tang, Jiang, *et al.*, 2013). For the MTG, other rs-fMRI studies reported increased functional activity in adults with antisocial personality disorder (Tang, Liu, *et al.*, 2013) and CU-related decreased rsFC in prison inmates (Espinoza *et al.*, 2018). Cerebellar and opercular regions have been implicated in CU-related rsFC alterations in adults (Philippi *et al.*, 2015) and juveniles (Aghajani *et al.*, 2016) and left insula in CU-related rsFC (Werhahn *et al.*, 2018), emotions (Lindquist *et al.*, 2012), and empathy (Lockwood *et al.*, 2013). The left MTG seems engaged in theory of mind (Bzdok *et al.*, 2012) and CU level-dependent fear response (Sebastian *et al.*, 2014). ITG and MTG are thought to be implicated in moral (Boccia *et al.*, 2017), emotions (Alegria, Radua and Rubia, 2016; Anderson *et al.*, 2017), and reward-related processes (Crowley *et al.*, 2010) along with bilateral cerebellum (Veroude *et al.*, 2016). Although our results did not show an overlap between CU traits and RA or PA symptoms-related rsFC, cases showed a positive correlation between CU scores and PA scores on a behavioral level, which points to a behavioral finding (Pihet *et al.*, 2015). In sum, our subtype-specific patterns support a careful assessment of the individual manifestations of aggression. This could result in tailoring therapeutic approaches (Umbach, Berryessa and Raine, 2015; Wilkinson, Waller and Viding, 2016). Given the demonstrated subtype-specific rsFC alterations neurobiological interventions such as real-time fMRI or arousal-biofeedback trainings seem promising.

### Study Strengths and Limitations

Strengths of our multi-center study include the large sample, recruitment of both males and females, children and adolescents, and not restricting inclusion to CD given that CU traits are present in only a minority of children with CD (Frick, 2016). Moreover, we focused on RA and PA symptoms and CU traits, applied unrestricted whole-brain analysis of rsFC, and controlled for ADHD and anxiety symptoms. Therefore, our results might enable conclusions on aggression subtypes of aggressive youths representing clinical practice. Among study limitations is the reduced data homogeneity due to data collection at different sites, which may have limited our study power. Yet, the large sample size and the multi-center design increased reliability and generalizability of our results. Further, we added site as dummy-coded covariate, as the distribution of cases and controls across sites was not balanced. Moreover, while 38 cases without a diagnosis exhibited lower PA scores, subsequent sensitivity analysis yielded comparable aggression subtype-specific associations.

The current study clarifies that distinct though overlapping brain connectivity measures can disentangle RA and PA aggression subtypes and behavioral modifiers, and that considering comorbid symptoms is crucial. The observed effects of controlling for both ADHD and anxiety on the rsFC in aggressive children and youths are even more extensive than previously thought. The partly overlapping associations for RA compared to PA symptoms and the distinct CU traits-related connectivities corroborate behavioral and neural findings. Affected brain areas have been previously implicated in processes including emotion and moral-related behaviors. Diagnostic and treatment approaches, however, largely disregarded particularly RA and PA forms of aggressive behavior. Investigating task-based alterations for these aggression subtypes could further widen the knowledge of underlying neural correlates and encourage the transfer of aggression subtype-specific procedures to the clinical practice.

## Supporting information

Supplement_2ndManuscript_Werhahn

## ACKNOWLEDGMENTS

This project has received funding from the European Union’s Seventh Framework Programme for research, technological development and demonstration under grant agreement 602805 (Aggressotype) and no 603016 (MATRICS). This manuscript reflects only the author’s view and the European Union is not liable for any use that may be made of the information contained herein. The authors express their deepest gratitude to all participating families.

Dr. Michael Craig is currently funded by the Medical Research Council UK (grant MR/M013588). Dr. Tobias Banaschewski served in an advisory or consultancy role for Actelion, Hexal Pharma, Lilly, Lundbeck, Medice, Neurim Pharmaceuticals, Novartis, Shire, received conference support or speaker’s fee by Lilly, Medice, Novartis and Shire, has been involved in clinical trials conducted by Shire & Viforpharma, and received royalties from Hogrefe, Kohlhammer, CIP Medien, Oxford University Press. Dr. Celso Arango has been a consultant to or has received honoraria or grants from Acadia, Ambrosseti, Gedeon Richter, Janssen Cilag, Lundbeck, Merck, Otsuka, Roche, Servier, Shire, Schering Plough, Sumitomo Dainippon Pharma, Sunovion and Takeda. Dr. Daniel Brandies serves as an unpaid scientific advisor for an EU-funded neurofeedback trial unrelated to the present work. Dr. Barbara Franke receives funding from a personal Vici grant of the Dutch Organisation for Scientific Research (grant 016 130 669) and from the Dutch National Science Agenda for the NWANeurolabNL project (grant 400 17 602), and received educational speaking fees from Shire and Medicine. Dr. Susanne Walitza received royalties from Thieme, Hogrefe, Kohlhammer, Springer, Beltz in the last five years. Her work in the last five years was supported by the Swiss National Science Foundation (SNF), diff. EU FP7s, HSM Hochspezialisierte Medizin of the Kanton Zurich, Switzerland, Bfarm Germany, ZInFP, Hartmann Müller Stiftung, Olga Mayenfisch Gertrud Thalmann Fonds. Outside professional activities and interests are declared under the link www.uzh.ch/prof/ssl-dir/interessenbindungen/client/web/. Dr. Jan Buitelaar has been in the past 3 years a consultant to/member of advisory board of/and/or speaker for Shire, Roche, Medice, and Servier. He is not an employee of these companies, and not a stock shareholder of any of these companies, and has no other financial or material support, including expert testimony, patents, and royalties. The present work is unrelated to the above grants and relationships. The authors do not have potential conflicts of interest.

