## Supplement_2ndManuscript_Werhahn for "Different Whole-Brain Functional Connectivity Correlates of Reactive-Proactive Aggression and Callous-Unemotional Traits in Disruptive Children and Adolescents"

**Supplementary Materials and Methods**

**Participants**

One hundred eighteen children and adolescents with a DSM-diagnosis of CD and/or ODD and/or aggression scores in the clinical range along with 89 age and handedness-matched healthy controls were included in the present study. Recruitment took place at nine different sites in Europe at resident hospitals, ambulatories, and eligible (boarding) schools (Radboud University Medical Center and the Donders Center for Cognitive Neuroimaging, Nijmegen, The Netherlands; Department of Child and Adolescent Psychiatry, University Medical Centre Groningen, The Netherlands; Central Institute of Mental Health, Medical Faculty, Mannheim/Heidelberg University, Mannheim, Germany; University of Ulm, Department of Child and Adolescent Psychiatry/Psychotherapy and Department of Psychiatry III, Ulm, Germany; Department of Child Psychiatry, and the Centre for Neuroimaging Sciences, Institute of Psychiatry, Psychology and Neuroscience, King`s College London, London, England; Institut d'Investigacions Biomèdiques August Pi i Sunyer and Hospital Clinic de Barcelona, Barcelona, Spain; Instituto de Investigación Sanitaria Gregorio Marañón, Child and Adolescent Psychiatry Department of Gregorio Marañòn General University Hospital, Madrid, Spain; Department of Child and Adolescent Psychiatry and Psychotherapy, University Zurich and MR center, Psychiatric University Hospital, Zurich, Switzerland; IRCCS Santa Lucia Foundation, Rome, Italy).

Besides a DSM-diagnosis of CD, ODD, and/or an aggression or rule-breaking behavior subscale score in a clinical range (T > 70) according to the CBCL, Youth Self Report (YSR), or Teacher Report Form (TRF), further inclusion criteria for the case group were no medication or a stable medication at least for two months. Exclusion criteria for cases was a primary DSM-diagnosis of depression, anxiety, psychosis, or bipolar disorder, and for the typically developed comparison group a DSM-diagnosis or clinically relevant scores in the CBCL, YSR, or TRF. Further exclusion criteria for all participants were an IQ score < 80 as measured by the Wechsler Intelligence Scale for Children, Fourth Edition (WISC-IV), and common contraindications for MRI scanning, such as braces or metal parts. Additionally, an anxiety score >8 on a Visual Analogue Scale ranging from 1 to 10 before scanning led to exclusion, as anxiety caused by the scanner could impair data quality and alter neural activity (Muehlhan, Lueken, Wittchen, & Kirschbaum, 2011). Participants had sufficient native language skills according to the assessing country. All sites obtained ethical approval separately. After participants received information about the study procedure, participants and their parents or legal representatives gave written informed consent.

**Assessment Tools and Study Procedure**

Diagnostic assessments and magnetic resonance (MR) measurements took place on different dates to minimize burden for the participants. During the first appointment, parents/primary caregivers and children/youths were assessed by trained (clinical) psychologists or accordingly instructed interns separately with the semi-structured interview Kiddie-Schedule for Affective Disorders and Schizophrenia, present and lifetime version K-SADS (Kaufman et al., 1997). The full supplementary module of the specific disorder followed positively answered questions. Diagnoses resulted from clinical evaluation after self- and parent-reports. If not already completed and brought to the assessment, parents/primary caregivers and children/youths answered a questionnaire package that included the following measures. The Child Behavior Checklist (CBCL) is a parent-report questionnaire rating the child or adolescent on various behavioral and emotional problems (Achenbach, 1991). The Swanson, Nolan, and Pelham teacher and parent rating scale (SNAP-IV) contains 26 items to measure attention deficit disorder (ADHD) and oppositional defiant disorder (ODD) symptoms from childhood to young adulthood (Swanson, 1992). The Inventory of Callous-Unemotional traits (ICU) is a 24-item questionnaire ranging from 0 (not at all true) to 3 (definitely true) (Frick, 2004). The Reactive-Proactive Aggression Questionnaire (RPQ) is a 23-item self-report measure of the frequency (never: 0, sometimes: 1, often: 2) of reactive (12 items) and proactive aggression (11 items) (Raine et al., 2006). IQ was estimated according to four sub-tests derived from the Wechsler Intelligence Scale for Children (WISC-IV) (Wechsler, 2003): block design, similarities, vocabulary, picture completion. Additionally, digit span subtest was assessed. Further parts of test battery in the framework of the Aggressotype/MATRICS study were also conducted as included additional questionnaires (Modified Aggression Scale (Kay, Wolkenfeld, & Murrill, 1988), Longterm difficulties self- and parent-report (Ormel et al., 2012), Pubertal Developmental Scale (Petersen, Crockett, Richards, & Boxer, 1988), Antisocial Behavior Scale (Slot, Orobio De Castro, & Duivenvoorden, 2000), Strengths and Difficulties Questionnaire (Goodman, 1997), Youth Self Report (YSR) and Teacher Report Form (TRF) as further parts of Achenbach System of Empirically Based Assessment (Achenbach, 1991) as well as ICU (Frick, 2004) self- and parent-report, cognitive tests (Probabilistic Reversal Learning Task (Cools, Clark, Owen, & Robbins, 2002) and three tasks from the Cambridge Neuropsychological Test Automated Battery (Cognition, 1996): Emotion Recognition Task, Delayed Matching to Sample and Rapid Visual Information Sampling). Blood- or saliva-samples were used for biological analysis. Two sites additionally conducted a short psychophysiological measurement (including measures of heart rate (pulse), skin conductance level, and EEG before, during, and after an emotional paradigm). All children/youths were prepared for MR scanning in attendance of their parents/primary caregivers. They were presented with MR sounds, and visited a dummy scanner and/or watched a MR information video to become familiar with the scanner environment. If the participants or parent/primary caregiver reported an anxiety rating of >8 on a Visual Analogue Scale (VAS) ranging from 1 (no anxiety) to 10 (very high anxiety) related to entering the MR scanner, the participant was then excluded from further investigation.

If all inclusion criteria were fulfilled, MR scanning took place on another appointment. Participants answered a MR safety form and a short pre-scanning questionnaire (e.g., regarding tobacco and cannabis consummation, genetic diseases in the family) and then practiced the following fMRI tasks on a laptop: passive avoidance task (Finger et al., 2011), emotional matching task (Via et al., 2014), and stop signal task (Rubia, Halari, Mohammad, Taylor, & Brammer, 2011). Participants gave a saliva sample right before MRI scanning. They were asked to use the toilet, and female pregnancy was ruled out by a urine pregnancy test or based on self-reports. The first part of MRI scanning session included a T1-weighted anatomical image, the three fMRI tasks mentioned above, and the fMRI resting state sequence. For the resting state scan, participants were instructed to look at a white crosshair against a black background and let their mind wander. After a break that included the collection of another saliva sample and the assessment of a short questionnaire regarding the performance in the fMRI tasks, a T1-weighted anatomical scan was followed by two sequences of magnetic resonance spectroscopy and diffusion tensor imaging. After all study-related procedures, the granted monetary reward and travel costs were reimbursed. Additionally, participants received a picture of their anatomical MR scan. The current study reports resting state fMRI and psychometric data.

**Image Acquisition**

Supplementary Table 1 and Table 2 show the site-specific scanner information.

| **Supplementary Table 1.**  Structural MRI scan parameters across sites. | | | | | | | |
| --- | --- | --- | --- | --- | --- | --- | --- |
| **Scanner** | **Site** | **TR/TE/T1 (ms)** | **Flip angle** | **Field of view** | **Matrix RL/AP/slices** | **Voxel size (mm)** | **Acceleration factor** |
| Siemens | Nijmegen | 2300/2.98/900 | 9 | 256 | 212/256/176 | 1.0x1.0x1.2 | 2 |
|  | Mannheim | 2300/2.96/900 | 9 | 256 | 212/256/176 | 1.0x1.0x1.2 | 2 |
|  | Ulm | 2300/2.96/900 | 9 | 256 | 212/256/176 | 1.0x1.0x1.2 | 2 |
|  | Barcelona | 2300/2.98/900 | 9 | 256 | 212/256/176 | 1.0x1.0x1.2 | 2 |
|  | Madrid | 2300/2.98/900 | 9 | 256 | 212/256/176 | 1.0x1.0x1.2 | 2 |
|  | Rome | 2080/2.86/900 | 9 | 256 | 212/256/176 | 1.0x1.0x1.2 | 2 |
| Philips | Groningen | 2450/3.11/900 | 8 | 270 | 256/232/170 | 1.0x1.0x1.0 | 1.8 |
|  | Zurich | 2300/3.11/900 | 9 | 270 | 256/232/170 | 1.0x1.0x1.0 | 1.8 |
| GE | London | 2300/3.02/400 | 11 | 270 | 256/256/196 | 1.0x1.0x1.2 | 1.75 |

| **Supplementary Table 2.**  Resting state functional MRI scan parameters across sites. | | | | | | |
| --- | --- | --- | --- | --- | --- | --- |
| **Scanner** | **Site** | **TR, TE1/TE2/TE3 (ms)** | **Number of slices** | **Slice scan order** | **Voxel size (mm)** | **Duration (min)** |
| Siemens | Nijmegen | 2300, 12/28.4/44.8 | 33 | descending | 3.8x3.8x3.8 | 8:24 |
|  | Mannheim | 2300, 12/29/46 | 33 | descending | 3.8x3.8x3.8 | 8:24 |
|  | Ulm | 2300, 31 | 33 | descending | 3.8x3.8x3.8 | 8:23 |
|  | Barcelona | 2300, 12 | 33 | descending | 3.8x3.8x3.8 | 8:21 |
|  | Madrid | 2300, 13 | 36 | descending | 3.8x3.8x3.8 | 8:24 |
|  | Rome | 2080, 30 | 32 | ascending | 3.0x3.0x2.5 | 7:38 |
| Philips | Groningen | 2450, 8.01/22.02/36.02 | 45 | descending | 3.5x3.5x3.5 | 10:08 |
|  | Zurich | 2300, 13/31/49 | 33 | descending | 3.75x3.75x3.79 | 7:51 |
| GE | London | 2300, 11.8/31/48 | 33 | descending, interleaved | 3.45x3.45x4.20 | 8:15 |

**Supplementary Results**

The distribution of demographic characteristics, diagnoses, and clinical aggression scores on behavioral measures across sites are shown in Supplementary Table 3. The investigated behavioral measures exhibited considerable standard deviations, which enabled the planned dimensional approach of aggression dimension-specific rsFC analyses.

In the whole sample, RA and PA symptoms correlated positively with ADHD inattention subscale (*r* = 0.50 and *r* = 0.34, *p* < 0.001) and ADHD hyperactivity/impulsivity subscale (*r* = 0.50 and *r* = 0.43, *p* < 0.001) and with anxiety problems (*r* = 0.36, *p* < 0.001 and *r* = 0.19, *p* = 0.02). CU traits also related positively to ADHD inattention and hyperactivity/impulsivity subscales (*r* = 0.59 and *r* = 0.49, *p* < 0.001) and to anxiety levels, yet not reaching significance (*r* = 0.08, *p* > 0.05). Within cases, RA and PA symptoms did not correlate with ADHD inattention subscale (*r* = -0.01, *p* > 0.05 and *r* = -0.04, *p* > 0.05) nor hyperactivity/impulsivity subscale (*r* < 0.01, *p* > 0.05 and *r* = 0.12, *p* > 0.05). Only the positive correlation of RA (*r* = 0.32, *p* = 0.004) but not PA (*r* = 0.10, *p* > 0.05) with anxiety symptoms yielded significance. CU traits were positively associated with ADHD inattention subscale (*r* = 0.32, *p* = 0.001) but with neither hyperactivity/impulsivity subscale (*r* = 0.13, *p* > 0.05) nor anxiety problems (*r* = -0.06, *p* > 0.05).

Cases showed positive correlations between RA and PA (*r* = 0.63, *p* < 0.001) and between CU traits and PA (*r* = 0.33, *p* = 0.001), while the positive association of CU traits and RA did not reach significance (*r* = 0.13, *p* > 0.05).

The average RMS-FD for cases was 0.12mm (*SD* = 0.17mm) and for controls 0.09 (*SD* = 0.18mm). In the whole sample, mean RMS-FD correlates positively with CU traits (*r* = 0.17, *p* < 0.05), while associations with CBCL rule-breaking and aggression subscales, reactive and proactive scores along with ADHD inattention and hyperactivity/impulsivity subscales and anxiety problems did not reach significance (*p* > 0.05) or trend level. Within cases, there was no significant or trend level correlation between mean RMS-FD and these clinical characteristics. As cases and controls differed in average RMS-FD values, we conducted sensitivity analyses for our case-control differences in voxel-wise rsFC (when we additionally controlled for anxiety but not ADHD symptoms) and additionally included RMS-FD as covariate. Compared to controls, cases showed comparable decreased ICC in a cluster including the right occipital pole (*T*(141) = -5.28, *p*-FDR < 0.05) and increased ILC in a left hemispheric frontal cluster including bilateral frontal pole extending to the medial frontal cortex (*T*(141) = 6.23, *p*-FDR < 0.05).

For all voxel-to-voxel whole-brain rsFC analysis, a statistical threshold of *p* < 0.001, *p* < 0.05 FDR cluster-level correction for multiple comparisons was applied (see Supplementary Table 4 and 5). After primary analyses that included the control for site, we additionally controlled for age, sex, IQ, medication, and handedness subsequently, as they can possibly influence rsFC of RSNs such as the DMN (Allen et al., 2011; Mak et al., 2017; Sherman et al., 2014) or functional activity (Liu, Stufflebeam, Sepulcre, Hedden, & Buckner, 2009) (see Supplementary Table 4 and 5).

Regarding the observed case-control differences in rsFC (when controlled for anxiety levels besides site), sensitivity analyses within CONN showed that the reduced ICC in an right occipital cluster in cases compared to control only survived the additional post hoc control for age and sex (*T*(140) = -5.35, *p*-FDR < 0.05). Furthermore, the increased ILC in a frontal cluster in cases compared to controls only survived post hoc control for age, sex, IQ, and medication but not handedness (*T*(131) = 5.67, *p*-FDR < 0.05).

Given the observed effects of the control for both ADHD and anxiety symptoms on aggression-related rsFC, we further explored their influence separately on rsFC within cases. Anxiety problems were associated negatively with ICC in a right hemispheric cluster including temporal pole an anterior divisions of superior and middle temporal gyrus, when we controlled for site and ADHD symptoms. Moreover, only the ADHD inattention but not hyperactivity subscale related positively to ICC in a cluster including right superior lateral occipital cortex and positively to ILC in right hemispheric lingual gyrus, occipital fusiform gyrus, and cerebellum 6 regions, yet only when we controlled for site but not anxiety symptoms (all *p*-FDR < 0.05).

Furthermore, we explored rsFC patterns when not controlling for neither ADHD nor anxiety symptoms. We found increased ILC in a cluster including bilateral frontal pole only for cases compared to controls. Within cases, CU traits related to decreased ICC in a left hemispheric cluster including MTG and inferior lateral occipital cortex (all *p*-FDR < 0.05).

After taking ADHD symptoms but not anxiety problems into account, only one cluster in left central gyrus with proactive aggression-related reduced ILC survived. Another cluster extending from left hemispheric Heschl`s gyrus to superior temporal gyrus exhibited increased ILC with higher levels of CU-traits. When anxiety problems but not ADHD symptoms were controlled for, the majority of aggression subtype-specific rsFC patterns disappeared. Yet, the reactive aggression-specific decrease in correlation strength survived in one cluster including the left superior division of the parietal lobe. Furthermore, one CU traits-related left hemispheric cluster in ITG, MTG, and inferior lateral occipital cortex survived, showing decreased ICC with higher levels of CU traits (all *p*-FDR < 0.05).

Within cases, sensitivity analyses showed negative associations between RA aggression and ICC in two clusters (*r* = -0.49, *p* < 0.001, and *r* = -0.49, *p* < 0.001), which were also present after exclusion of 38 cases without a DSM-diagnosis of CD and/or ODD (*r* = -0.57, *p* < 0.001, and *r* = -0.60, *p* < 0.001). The positive PA symptoms-related increased ICC in another cluster (*r* = 0.40, *p* = 0.001) remained as well after their exclusion (*r* = 0.48, *p* < 0.001). Furthermore, the negative correlation between RA behavior and ILC (*r* = -0.40, *p* = 0.001) remained significant (*r* = -0.41, *p* = 0.002). Lastly, we found the significant negative associations of PA behavior and ILC in three clusters (*r* = -0.55, *p* < 0.001 and *r* = -0.40, *p* = 0.001 and *r* = -0.39, *p* = 0.001) also after the exclusion of cases without a diagnosis (*r* = -0.58, *p* < 0.001 and *r* = -0.41, *p* = 0.003 and *r* = -0.40, *p* = 0.003).

| **Supplementary Table 3.**  Distribution of demographic characteristics, diagnoses and aggression scores across sites. | | | | |
| --- | --- | --- | --- | --- |
|  |  | Cases (*N* = 118) | HC (*N* = 89) |  |
| Nijmegen (*N* = 40) | Age | 13.55 ± 2.59 | 12.64 ± 1.96 | 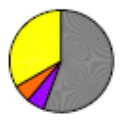 |
|  | IQ | 100.41 ± 11.83 | 107.86 ± 12.45 |  |
|  | Sex, m/f | 14/4 | 14/8 |  |
|  | Medication | 10 | 0 |  |
| Groningen  (*N* = 16) | Age | 14.77 ± 2.54 | 12.39 ± 2.32 | 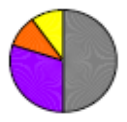 |
|  | IQ | 101.40 ± 11.63 | 101.85 ± 2.58 |  |
|  | Sex, m/f | 8/2 | 2/4 |  |
|  | Medication | 6 | 0 |  |
| Mannheim  (*N* = 33) | Age | 12.78 ± 2.42 | 12.97 ± 3.06 | 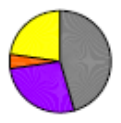 |
|  | IQ | 102.16 ± 10.13 | 116.32 ± 7.73 |  |
|  | Sex, m/f | 19/3 | 9/2 |  |
|  | Medication | 14 | 0 |  |
| Ulm (*N* = 18) | Age | 10.55 ± 2.38 | 13.59 ± 3.26 | 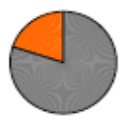 |
|  | IQ | 103.62 ± 17.55 | 101.41 ± 7.97 |  |
|  | Sex, m/f | 5/0 | 4/9 |  |
|  | Medication | 5 | 0 |  |
| London (*N* = 26) | Age | 14.67 ± 2.25 | 13.71 ± 2.05 | 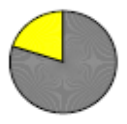 |
|  | IQ | 97.16 ± 10.33 | 111.85 ± 11.01 |  |
|  | Sex, m/f | 15/0 | 8/3 |  |
|  | Medication | 12 | 1 |  |
| Barcelona  (*N* = 23) | Age | 12.99 ± 2.82 | 14.94 ± 2.31 | 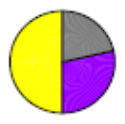 |
|  | IQ | 102.57 ± 12.10 | 105.70 ± 6.59 |  |
|  | Sex, m/f | 10/4 | 5/4 |  |
|  | Medication | 7 | 0 |  |
| Madrid (*N* = 20) | Age | 14.29 ± 2.22 | 14.68 ± 1.84 | 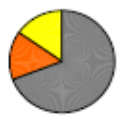 |
|  | IQ | 98.73 ± 9.45 | 105.74 ± 9.31 |  |
|  | Sex, m/f | 10/3 | 5/2 |  |
|  | Medication | 10 | 0 |  |
| Zürich  (*N* = 20) | Age | 10.53 ± 1.87 | 11.75 ± 1.49 | 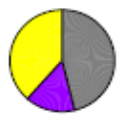 |
|  | IQ | 102.61 ± 11.20 | 99.60 ± 7.87 |  |
|  | Sex, m/f | 10/3 | 3/4 |  |
|  | Medication | 2 | 0 |  |
| Rome  (*N* = 11) | Age | 14.37 ± 2.60 | 16.12 ± 1.81 | 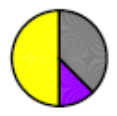 |
|  | IQ | 95.84 ± 8.83 | 97.23 ± 2.78 |  |
|  | Sex, m/f | 8/8 | 1/2 |  |
|  | Medication | 4 | 0 |  |
| 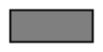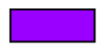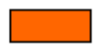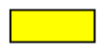*Note:* Values are means ± SD, or counts. HC = healthy controls. ODD CD ODD + CD     (additional) aggression in a clinical range (T > 70 on aggression subscales in Child Behavior Checklist)  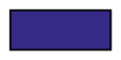Nijmegen 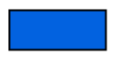 Groningen 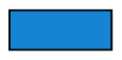Mannheim 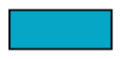 Ulm 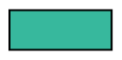 London 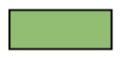 Barcelona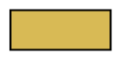 Madrid 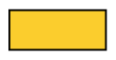 Zürich 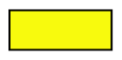 Rom  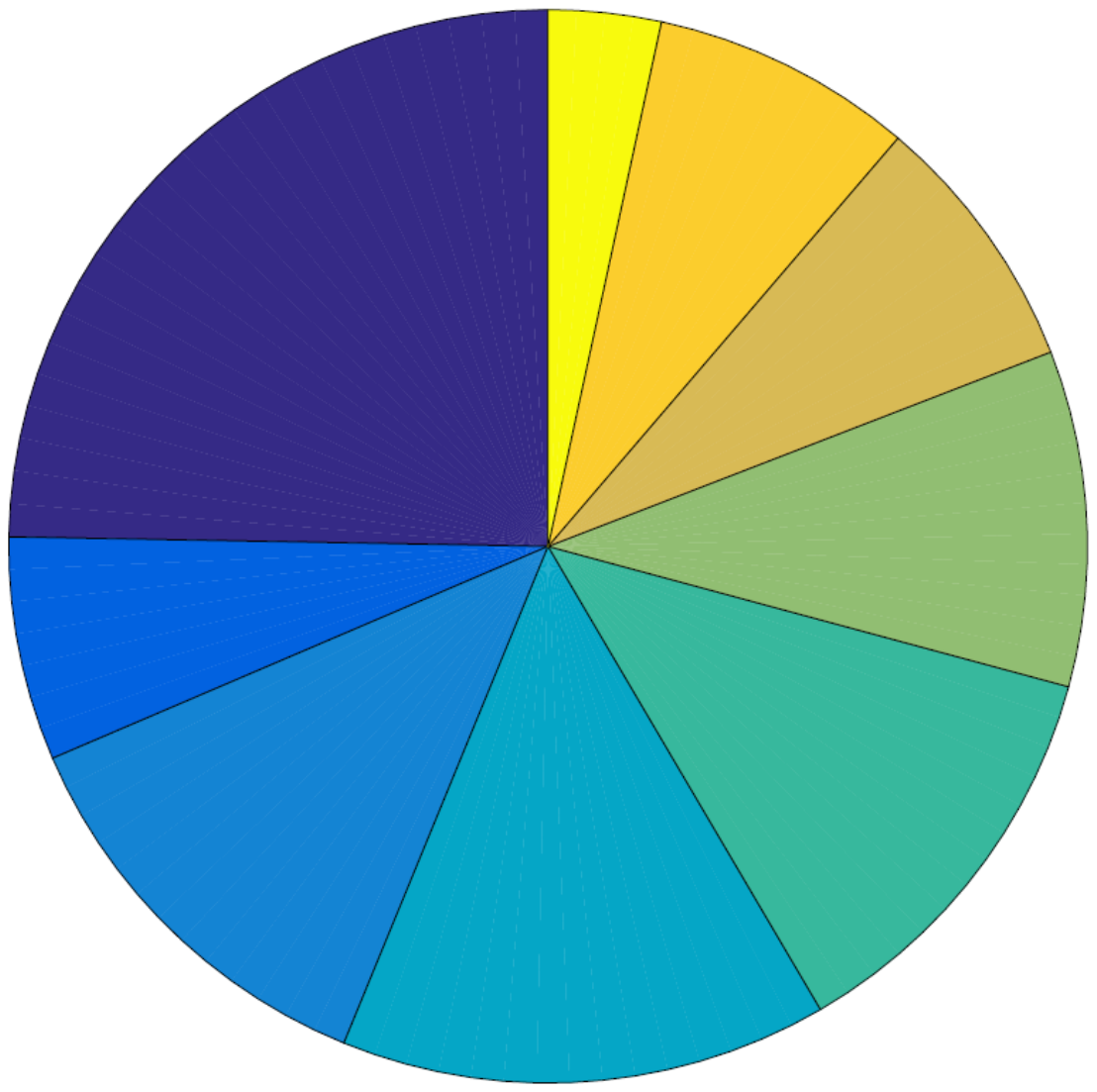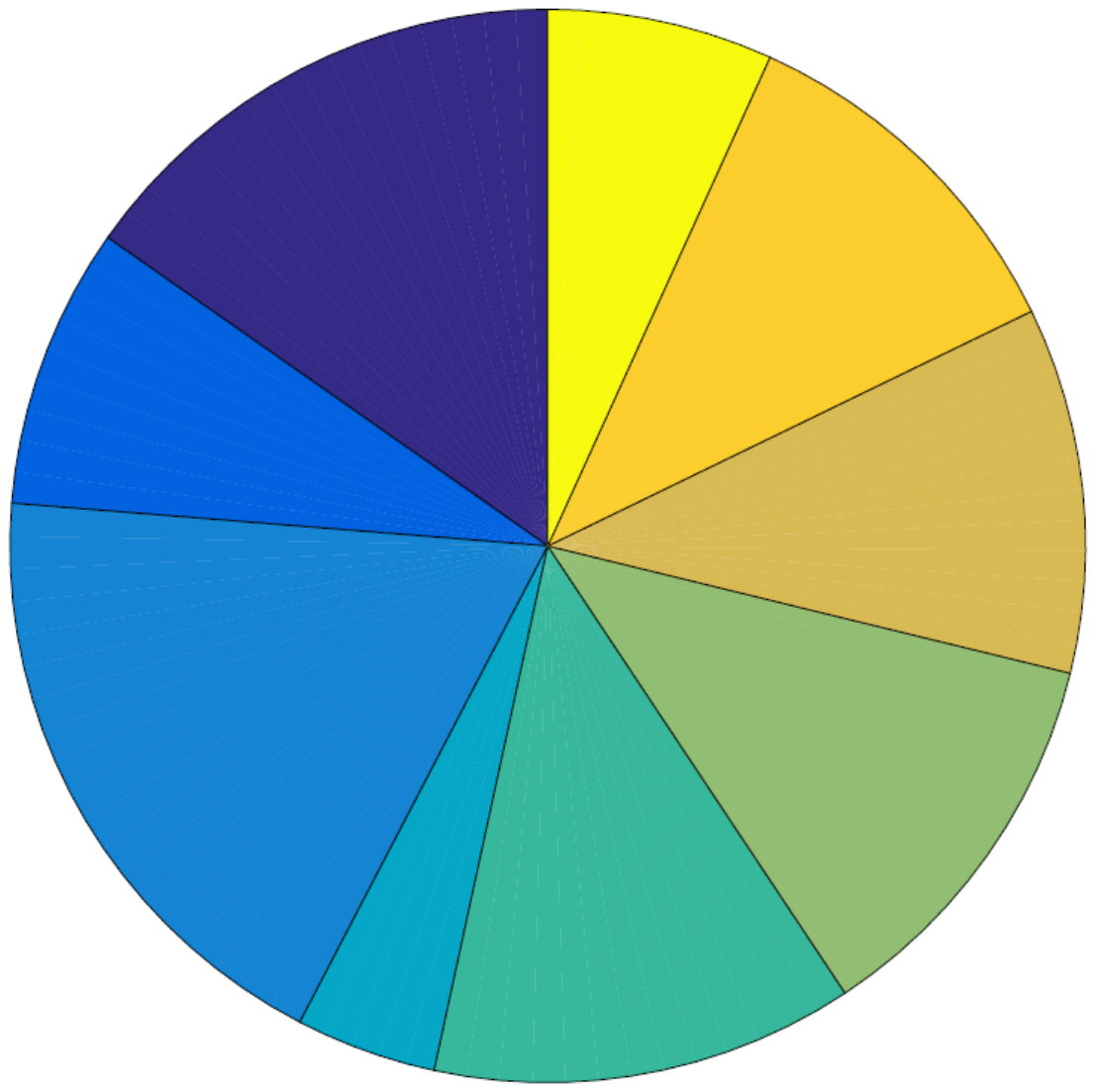 | | | | |

| **Supplementary Table 4.**  Clusters and coordinates derived from intrinsic connectivity contrast (ICC) results related to reactive and proactive aggression and callous-unemotional (CU) traits. | | | | | | | | | |
| --- | --- | --- | --- | --- | --- | --- | --- | --- | --- |
|  |  | | |  |  |  | Peak voxel  MNI coordinates | | |
|  | | Brain region | | Hemisphere | Cluster size | *β*-value | X | Y | Z |
| 1 | *Positive Association* - Inferior Temporal Gyrus  *Negative Association* - Inferior Temporal Gyrus  - Insula Lobe | | CU traits | L  L L | 130  139 59 | 0.38  -0.41 -0.36 | -54  -48 -40 | -52  -60 16 | -20  00 08 |
|  | *Negative Association* - Superior Parietal Lobe - Postcentral Gyrus | | Reactive Aggression | L L | 117 61 | -0.48 -0.49 | -32 -48 | -56 -12 | 68 58 |
|  | *Positive Association* - Superior Parietal Lobe | | Proactive Aggression | R | 88 | 0.27 | 26 | -56 | 62 |
| 2 | *Negative Association* - Inferior Temporal Gyrus - Insula Lobe | | CU traits | L L | 132 67 | -0.41 -0.35 | -48 -40 | -60 16 | 00 08 |
|  | *Positive Association* - Superior Parietal Lobe | | Proactive Aggression | R | 94 | 0.31 | 26 | -56 | 62 |
| *Note:* Brain regions and cytoarchitectonic coordinates of peak voxels for callous-unemotional (CU) traits and reactive and proactive aggression are labeled based upon Anatomy toolbox for SPM using Montreal Neurological Space (MNI) anatomical coordinates. Statistical thresholds for the reported results are *p* < 0.001, false discovery rate (FDR) cluster-level corrected (*p* < 0.05) for multiple comparisons. The β-values represent z-standardized correlation coefficients. Regression analyses of aggression subtype-specific degree centrality are corrected for site, ADHD and anxiety scores (1) and for site, age, gender, IQ, medication, and handedness along with ADHD and anxiety scores (2), respectively. | | | | | | | | | |

| **Supplementary Table 5.**  Clusters and coordinates derived from integrated local correlation (ILC) analyses for reactive and proactive aggression. | | | | | | | | | |
| --- | --- | --- | --- | --- | --- | --- | --- | --- | --- |
|  |  | | |  |  |  | Peak voxel  MNI coordinates | | |
|  | | Brain region | | Hemisphere | Cluster size | *β*-value | X | Y | Z |
| 1 | *Negative Association* - Superior Parietal Lobe | | Reactive Aggression | L | 288 | -0.33 | -32 | -54 | 68 |
|  | *Negative Association* - Precentral Gyrus - Superior Parietal Lobe - Inferior Parietal Lobe | | Proactive Aggression | L L L | 101 85 83 | -0.28 -0.21 -0.22 | -42 -26 -44 | -16 -56 -48 | 62 70 60 |
| 2 | *Positive Association* - Superior Parietal Lobe | | Reactive Aggression | L | 187 | -0.36 | -38 | -50 | 64 |
|  | *Negative Association* - Inferior Parietal Lobe - Precentral Gyrus - Superior Frontal Gyrus - Precentral Gyrus | | Proactive Aggression | L L R L | 201 160 139 99 | -0.24 -0.30 -0.22 -0.26 | -44 -42 24 -60 | -48 -16 28 -06 | 60 62 60 36 |
| *Note:* Brain regions and cytoarchitectonic coordinates of the peak voxels are labeled based upon Anatomy toolbox for SPM using Montreal Neurological Space (MNI) anatomical coordinates. The statistical threshold for the reported results is *p* < 0.001, false discovery rate (FDR) cluster-level corrected (*p* < 0.05) for multiple comparisons. β-values represent z-standardized correlation coefficients. Regression analyses of local coherence are corrected for site, ADHD and anxiety scores (1) and for site, age, gender, IQ, medication, and handedness along with ADHD and anxiety scores (2), respectively. | | | | | | | | | |

| **Supplementary Table 6.**    Bivariate correlations of aggression-related scores within cases. | | | | | | | | | | |
| --- | --- | --- | --- | --- | --- | --- | --- | --- | --- | --- |
|  | rule-break-ing | aggres-sion | ODD | CD | inatten-tion | hyper-activity/ impul-sivity | anxiety | CU traits | reactive aggres-sion | pro-active aggres-sion |
| rule-break-ing |  | *r* = 0.47,  *p* < 0.001 | *r* = 0.16,   *p > 0.05* | *r* = 0.38, *p* < 0.001 | *r* = 0.19,  *p* > 0.05 | *r* = 0.20,  *p* = 0.05 | *r* = -0.02,  *p* > 0.05 | *r* = 0.16, *p* > 0.05 | *r* = 0.05,  *p* > 0.05 | *r* = 0.14, *p* > 0.05 |
| aggres-sion | *r* = 0.47, *p* < 0.001 |  | *r* = 0.20,  *p* < 0.05 | *r* = 0.21,  *p* < 0.05 | *r* = 0.22,  *p* < 0.05 | *r* = 0.26,  *p* < 0.01 | *r* = -0.09,  *p* > 0.05 | *r* = 0.34,  *p* < 0.001 | *r* = 0.16,  *p* > 0.05 | *r* = 0.22,  *p* < 0.05 |
| ODD | *r* = 0.16, *p* > 0.05 | *r* = 0.20,  *p* < 0.05 |  | *r* = 0.28,  *p* < 0.01 | *r* < -0.01,  *p* > 0.05 | *r* = 0.13,  *p* > 0.05 | *r* = -0.02,  *p* > 0.05 | *r* = 0.18,   *p > 0.05* | *r* = 0.07,  *p* > 0.05 | *r* = 0.22,  *p* < 0.05 |
| CD | *r* = 0.47, *p* < 0.001 | *r* = 0.21,  *p* < 0.05 | *r* = 0.28,  *p* < 0.01 |  | *r* = 0.14,  *p* > 0.05 | *r* = 0.01,  *p* > 0.05 | *r* = -0.02,  *p* > 0.05 | *r* = 0.26,  *p* < 0.01 | *r* = 0.13,  *p* > 0.05 | *r* = 0.29,  *p* < 0.01 |
| inatten-tion | *r* = 0.12, *p* > 0.05 | *r* = 0.12,  *p* > 0.05 | *r* = 0.33,  *p* < 0.001 | *r* = 0.10,  *p* > 0.05 |  | *r* = 0.55,  *p* < 0.001 | *r* = -0.16,  *p* > 0.05 | *r* < 0.01,  *p* > 0.05 | *r* = 0.11,  *p* > 0.05 | *r* = -0.12,  *p* > 0.05 |
| hyper-activi-ty/ impul-sivity | *r* = 0.02, *p > 0.05* | *r* = 0.09,  *p* > 0.05 | *r* = 0.28,  *p* < 0.01 | *r* = 0.05,  *p* > 0.05 | *r* = 0.55,  *p* < 0.001 |  | *r* = -0.14,  *p* > 0.05 | *r* = -0.05,  *p* > 0.05 | *r* = 0.06,  *p* > 0.05 | *r* = 0.01,  *p* > 0.05 |
| anx-iety | *r* = 0.04, *p* > 0.05 | *r* = 0.14,  *p* > 0.05 | *r* = 0.42,  *p* < 0.001 | *r* = 0.07,  *p* > 0.05 | *r* = -0.16,  *p* > 0.05 | *r* = -0.14,  *p* > 0.05 |  | *r* = -0.08,  *p* > 0.05 | *r* = 0.08,  *p* > 0.05 | *r* < 0.01,  *p* > 0.05 |
| CU traits | *r* = 0.16, *p* > 0.05 | *r* = 0.34,  *p* < 0.001 | *r* = 0.18,   *p > 0.05* | *r* = 0.26,  *p* < 0.01 | *r* = 0.32,  *p* = 0.01 | *r* = 0.13,  *p* > 0.05 | *r* = -0.06,  *p* > 0.05 |  | *r* = 0.13,  *p* > 0.05 | *r* = 0.33,  *p* = 0.001 |
| reac-tive  aggres-sion | *r* = 0.05, *p* > 0.05 | *r* = 0.16,  *p* > 0.05 | *r* = 0.07,  *p* > 0.05 | *r* = 0.13,  *p* > 0.05 | *r* = -0.01,  *p* > 0.05 | *r* < 0.01,  *p* > 0.05 | *r* = 0.32,  *p* < 0.01 | *r* = 0.13,  *p* > 0.05 |  | *r* = 0.63,  *p* < 0.001 |
| pro-active aggres-sion | *r* = 0.14, *p* > 0.05 | *r* = 0.22,  *p* < 0.05 | *r* = 0.22,  *p* < 0.05 | *r* = 0.29,  *p* < 0.01 | *r* = -0.04,  *p* > 0.05 | *r* = 0.12,  *p* > 0.05 | *r* = 0.10,  *p* > 0.05 | *r* = 0.33,  *p* = 0.001 | *r* = 0.63,  *p* < 0.001 |  |
| *Note:* Rule-breaking and aggression T-scores derived from the Child Behavior Checklist. ODD and CD scores according to the Kiddie-Schedule for Affective Disorders and Schizophrenia, present and lifetime version. Callous-unemotional (CU) traits, total score, derived from the parent-reported Inventory of Callous-Unemotional traits, reactive and proactive aggression scores according to the self-reported Reactive-Proactive Aggression Questionnaire. Inattention, hyperactivity/impulsivity scores according to the Swanson, Nolan, and Pelham teacher and parent rating scale (SNAP-IV), anxiety problems derived from the Youth Self Report. | | | | | | | | | | |
